# Eco-evo-devo implications and archaeobiological perspectives of wild and domesticated grapevines fruits covariating traits

**DOI:** 10.1101/513036

**Authors:** Vincent Bonhomme, Sandrine Picq, Sarah Ivorra, Allowen Evin, Thierry Pastor, Roberto Bacilieri, Thierry Lacombe, Isabel Figueiral, Jean-Frédéric Terral, Laurent Bouby

## Abstract

The phenotypic changes that occurred during the domestication and diversification of grapevine are well known, particularly changes in seed morphology, but the functional causes and consequences behind these variations are poorly understood. Wild and domesticate grapes differ, among others, in the form of their pips: wild grapes produce roundish pips with short stalks and cultivated varieties have more elongated pips with longer stalks. Such variations of form are of first importance for archaeobotany since the pip form is, most often, the only remaining information in archaeological settings. This study aims to enlight archaeobotanical record and grapevine pip development by better understanding how size and shape (co)variates between pip and berry in both wild and domesticated *Vitis vinifera*. The covariation of berry size, number of seeds per berry (“piposity”), pip size and pip shape were explored on 49 grapevine accessions sampled among Euro-Mediterranean traditional cultivars and wild grapevines. We show that for wild grapevine, the higher the piposity, the bigger the berry and the more elongated the pip. For both wild and domesticated grapevine, the longer is the pip, the more it has a “domesticated” shape. Consequences for archaeobotanical studies are tested and discussed, and these covariations allowed the inference of berry dimensions from archaeological pips from a Southern France Roman site. This systematic exploration sheds light on new aspects of pip-berry relationship, in both size and shape, on grapevine eco-evo-devo changes during domestication, and invites to explore further the functional ecology of grapevine pip and berry and notably the impact of cultivation practices and human selection on grapevine morphology.

## Introduction

Grapevine (*Vitis vinifera* L.) is one of the most cultivated fruit species in the world [1], and has held a central economic and cultural role since ancient times, particularly in the Mediterranean area (Brun, 2003; McGovern, 2007). The berries of grapevine are primarily used in wine production, but can be consumed fresh or dried (i.e. table grape). The wild progenitor of grapevine, *Vitis vinifera* subsp. *sylvestris*, was first domesticated in the South Caucasian area [4], which has yielded the oldest wine making evidence (McGovern et al., 2017) dated to early Neolithic period (∼8000 BP). The existence of other domestication centres has also been argued [6,7]. Since the early times of domestication, grapevine varieties (or cultivars or “cépages”) of *Vitis vinifera* subsp. *vinifera* have been selected and propagated; today there are several thousand varieties, identified by ampelography (i.e. grape morphology) and molecular markers [8,9]. *V. vinifera* subspecies differ in their reproductive biology and other phenotypic changes following domestication include larger bunches, larger berries, higher diversity in berry shape and skin colour, and higher sugar content [10,11].

The quantitative morphological description of archaeobotanical material has brought major insights into the intertwined relationships between humans and domesticated plants [12–20], including grapevine (Bouby et al., 2013; Karasik, Rahimi, David, Weiss, & Drori, 2018; Mangafa & Kotsakis, 1996; Orrù, Grillo, Lovicu, Venora, & Bacchetta, 2013; Pagnoux et al., 2015; Terral et al., 2010; Ucchesu et al., 2016). So far, molecular approaches on ancient grapevine have yielded limited information on domestication [28,29], with the study of ancient DNA hindered by its poor preservation in charred archaeobotanical material (but see Ramos-Madrigal et al., 2019).

Wild and domesticate grape seeds differ in their form (i.e. size plus shape): wild grapes produce roundish pips with short stalks and cultivated varieties produce more elongated pips with longer stalks [31]. Such form variations have been identified on archaeological grapevine pips (Bouby et al., 2013; Pagnoux et al., 2015; Terral et al., 2010). Archaeological material is often charred which can cause domesticated pips to appear more similar to wild pips [32] yet experimental charring has demonstrated the robustness of identification (Bouby et al., 2018; Ucchesu et al., 2016).

The functional causes and consequences, if any, behind the form variation of grapevine pip are still poorly understood. If size, shape, taste and colour of berries are phenotypic traits that have been selected by humans, pip shape was likely not a direct target of selective pressures but may possibly be affected by: the berry size; the number of pips per berry; the growing environment and cultivation practices; the domestication status and the variety for domesticated grapevine; and developmental stochasticity. For instance, previous works suggested that pip size and the number of pips per berry are positively correlated to berry size [8,34,35] and that this correlation is stronger for wild grapevines (Bouby et al., 2013).

Direct selection for numerous and larger berries is likely as they are key yield factors, but to what extent the form changes observed in archaeological pips imply changes in the form of berries? How this relation could be affected by the cultivation and subsequent domestication of wild individuals? Pips with a form similar to that of modern wild grapevine pips have been repeatedly found in ancient viticultural sites, which is suggestive of cultivated “true” wild grapevines or “weakly” domesticated forms alongside “true” fully domesticated varieties (Bouby et al., 2013; Pagnoux et al., 2015).

This paper scrutinizes how the form of berries and pips they contain covariate. A dataset of domesticated and wild contemporary grapevines allowed to compare patterns of covariation between these two, wild and domestic, *Vitis vinifera* compartments. This article is divided into four questions: i) how does size (co)vary between pips and berries, and depending on the number of pips?; ii) how does shape (co)vary between pips and berries and depending on the number of pips per berry?; iii) how much pip shape depends on berry size, number of pips per berry, status, accession, and which practical consequences for archaeobotanical studies? iv) can we infer the dimensions of the berry dimensions from the (recovered archaeological) pips?

## Results

### Preliminary analyses on modern material

The average piposity is equivalent between domesticated and wild accessions (mean±sd: domesticated=2.01±0.891, wild=2.1±0.968; generalized linear model with Poisson error: df=1468, z=1.234, P=0.217 – Figure 2). The distribution of piposity, however, does differ (Fisher’s exact test: P=0.004) due to a higher proportion of 4-pips berries in wild grapevines (Fisher’s exact test with 4-pips berries removed: P=0.6272). No difference was observed between cultivated (domesticated and wild) accessions and those collected from wild (mean±sd: cultivated=2.06±0.931, non-cultivated=2.00±0.902; generalized linear model with Poisson error: df=1468, z=-0.676, P=0.499).

**Figure 1.**
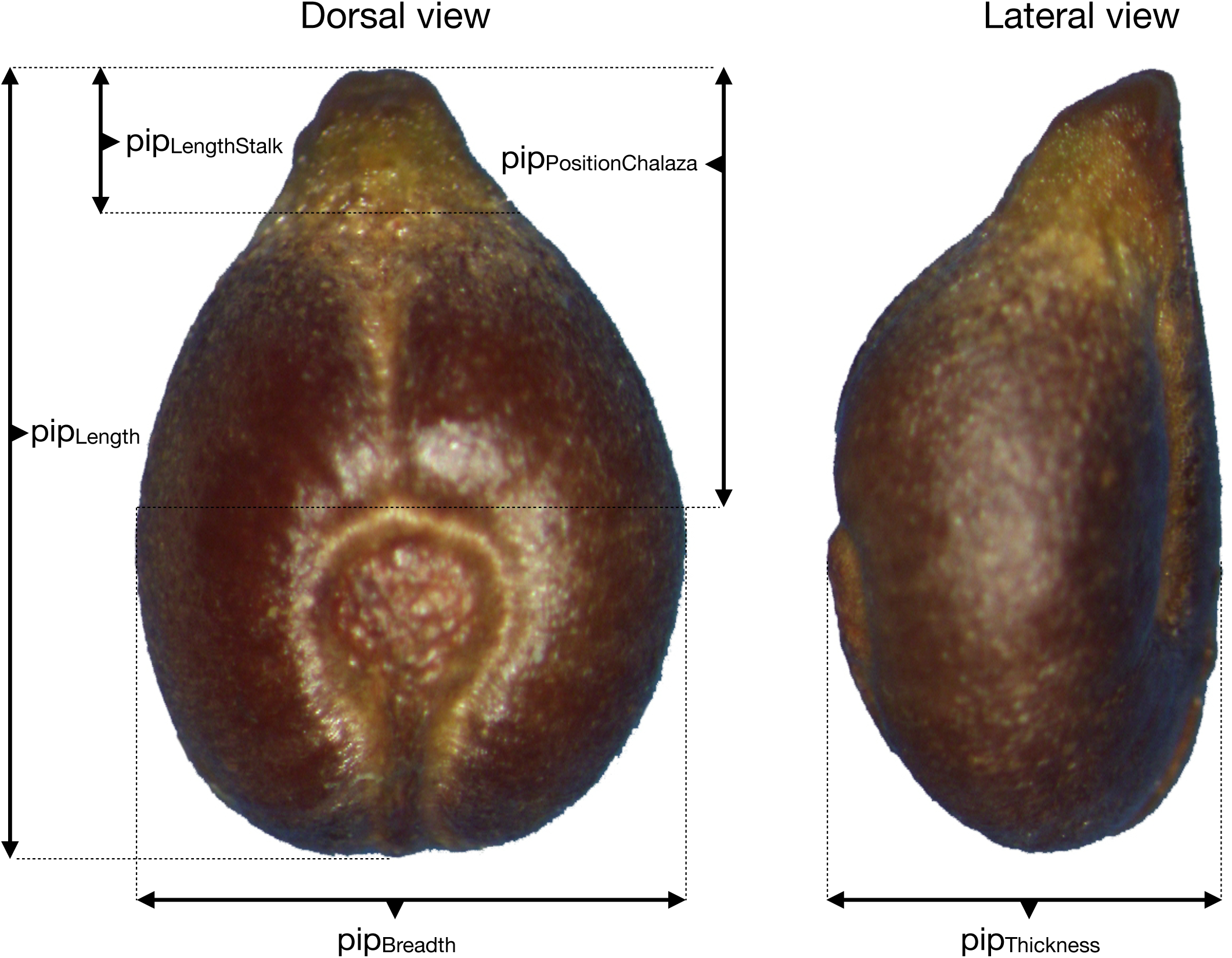
Dorsal and lateral views of a grapevine pip, here from the *Vitis vinifera* subsp. *sylvestris* wild” individual “Pic Saint-Loup 13”, with indications of morphometric measurements: pip_length_ (total length), pip_stalk_ (length of the stalk), pip_chalaza_ (position of the chalaza), pip_width_ and pip_thickness_.

**Figure 2.**
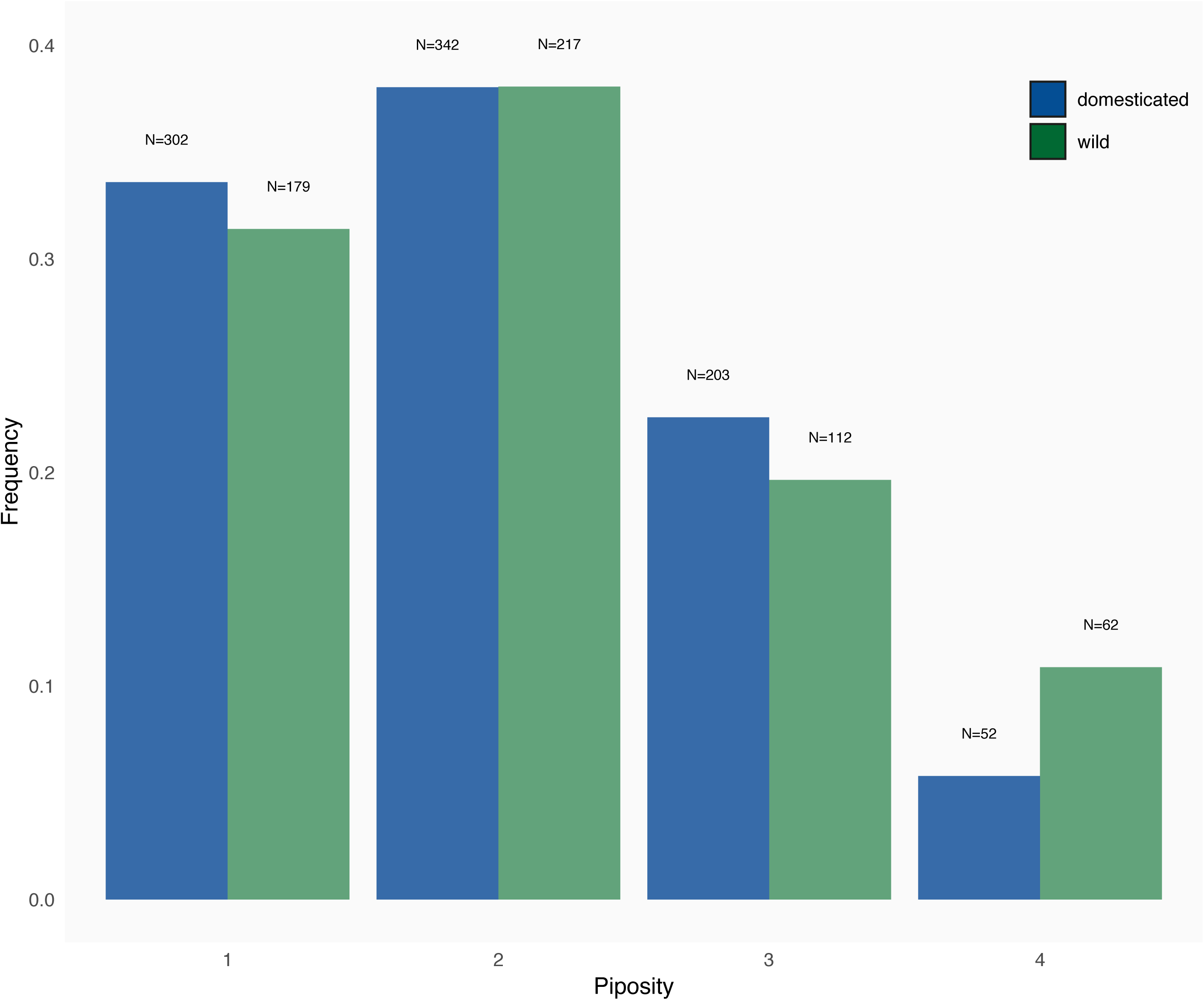
Distribution of the number of pips per berry for wild and domesticated grapevines.

### Covariation between pip and berry size in relation to the number of pips

#### Wild vs. domesticated

All berries and pips measurements were overall smaller for wild accessions compared to their domesticated counterparts (Wilcoxon one-tailed rank tests: all P<10^−8^ – Figure A). The extent of the difference between wild and domesticated varied between the pip dimensions (pip_LengthStalk_ > pip_PositionChalaza_ > pip_Length_ > pip_Thickness_ > pip_Breadth_).

Overall, the higher the piposity, the lower the contrast between domesticated and wild (Figure 3a). Pip dimensions of wild grapevines increase more substantially with increased piposity than their domesticated counterpart decrease. In wild grapevines, larger berries have more and larger pips (Figure 3a). No marked differences in berry dimensions/mass along increasing piposity were found for the cultivated grapevines, excepted between 1- and 2-pip for pip_Thickness_ (Wilcoxon rank tests: P=0.006).

**Figure 3.**
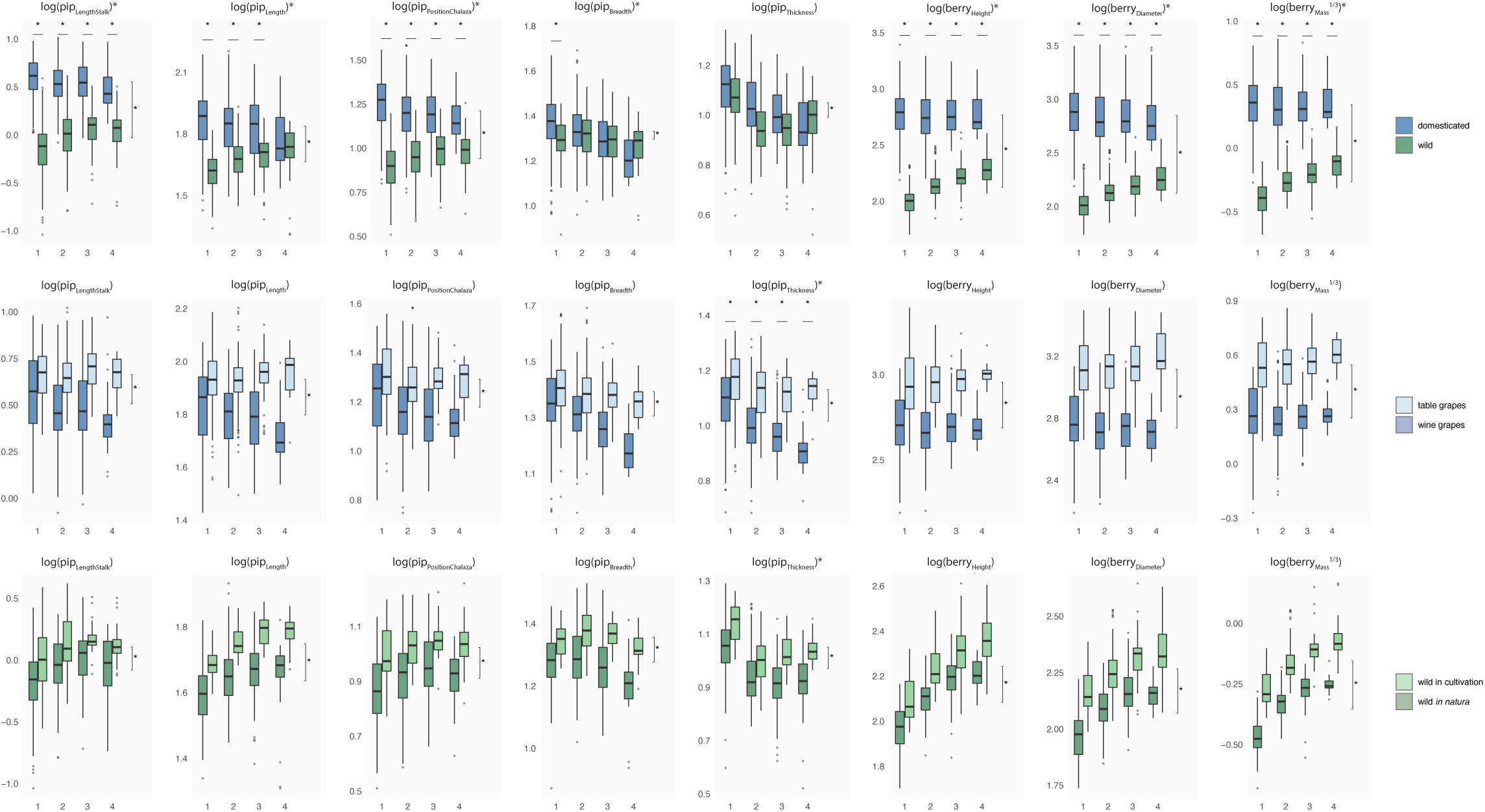
Comparisons of (logged) lengths and (log cubic-rooted) mass measurements. Each row represents a different comparison: a) wild and domesticated grapevines, b) table and wine varieties for domesticated accessions only, and c) cultivated and collected from wild for wild grapevine only. For each measurement, boxplots are displayed for each piposity level. Differences are tested using multivariate analyses of covariances, and differences of P<10^−5^, are indicated by stars in the facet title (interaction), on the right (overall difference) and above each piposity level (difference within a given piposity).

**Figure 4.**
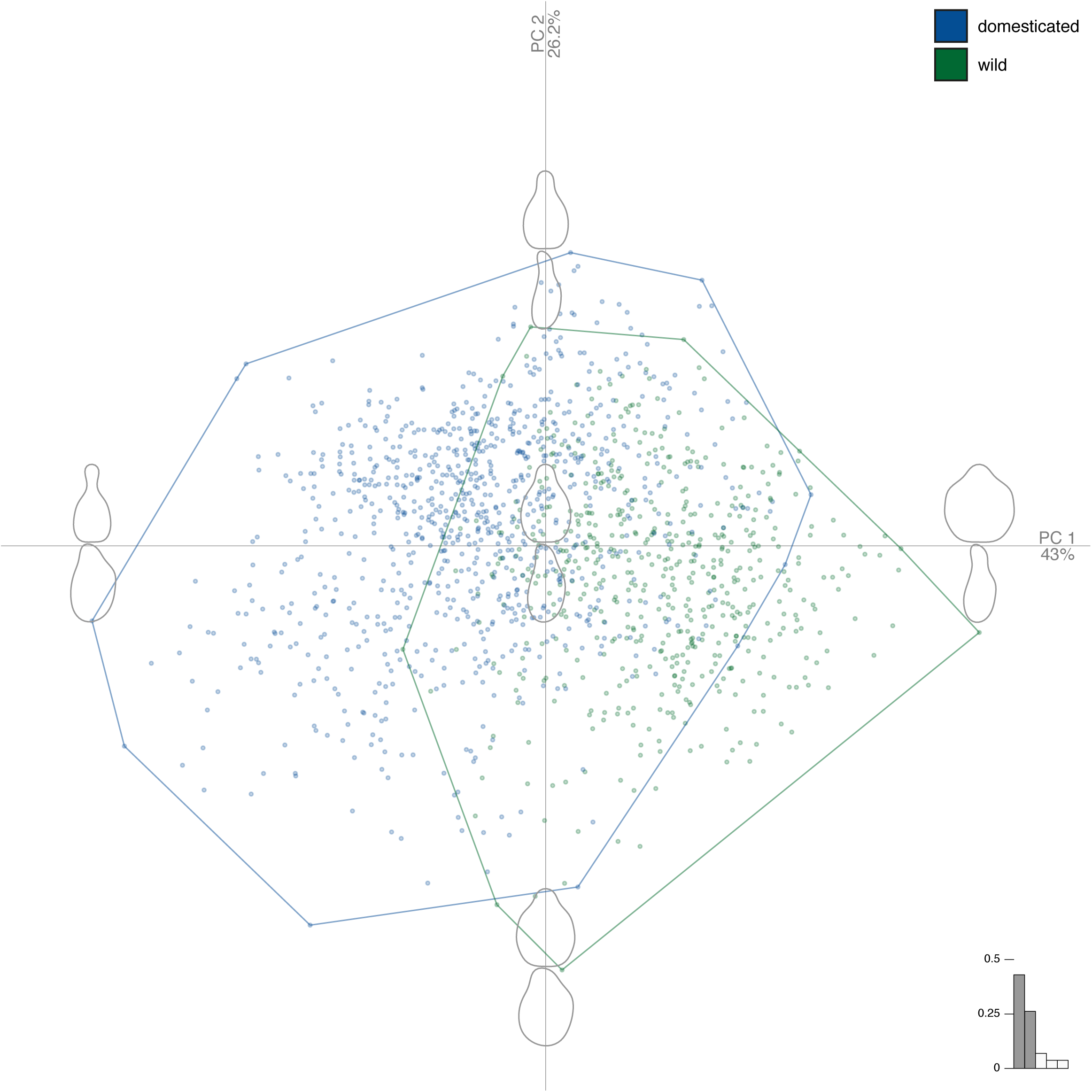
Principal component analysis on the joint matrices of Fourier coefficients obtained for the two views. The first two principal component gathered 69% of the total variance. The component of shape variation they capture are illustrated with reconstructed shapes at each extreme of their range. Colour of markers and convex hulls indicate pips from wild (green) and domesticated (blue) grapevines.

#### Table vs. wine cultivars

Table grapes have significantly higher dimensions than wine varieties (Figure 3b). With increasing piposity, table varieties tend to have bigger berries which is not the case for wine varieties. For pips, the only significant differences between low and high piposity were found for wine varieties and for pip_Breadth_ and pip_Thickness_ (P<10^−16^).

#### Wild grown in collection vs. wild in natura

Wild grapevine pips and berries are bigger when in cultivation than their counterparts growing *in natura* (Figure 3c). Besides these global differences, trends of all measured variables are similar along increasing piposity. The berry mass ratios, relatively to wild collected *in natura*, were on average, 6.4 for wine varieties, 15.6 for table ones and 1.8 for cultivated wild.

Bivariate comparisons (Figure B, Supplementary information) indicate positive correlations between all measurements. The total pip_Length_ appears to be the most consistent variable, between domesticated and wild grapevines: indeed, only the correlation with the pip_LengthStalk_ show a significant interaction. Inversely, the correlations implying pip_LengthStalk_ always show a significant interaction. For pips dimensions, the best correlations were found between pip_Length_ and pip_PositionChalaza_ (adj. r^2^=0.8) among those with non-significant interactions, and between pip_LengthStalk_ and pip_PositionChalaza_ (adj. r^2^_wild_=0.615, adj. r^2^_domesticated_=0.717) among those with significant interactions. Compared to pips dimensions, correlations between berry dimensions were much better and the three possible interactions were all significant.

### Covariation between pip and berry shape in relation to the number of pips

The PCA shows that the first two PCs (Figure 5) gathered 69% of the total shape variation, and higher rank components clearly levelled off (PC1=43.0%; PC2=26.2%; PC3=6.7%; PC4=3.9%), only the first two PCs were used as synthetic shape variables. Shape differences between wild and domesticated grapevines are mostly captured on PC1 yet scores on both PC1 (Wilcoxon rank tests, P<10^−16^) and on PC2 (P<10^−16^) were found different. Here, PC1 represents how prominent is the stalk and how round is the pip; PC2 represents the circularity, a more global length/width ratio of pips, for the two views.

**Figure 5.**
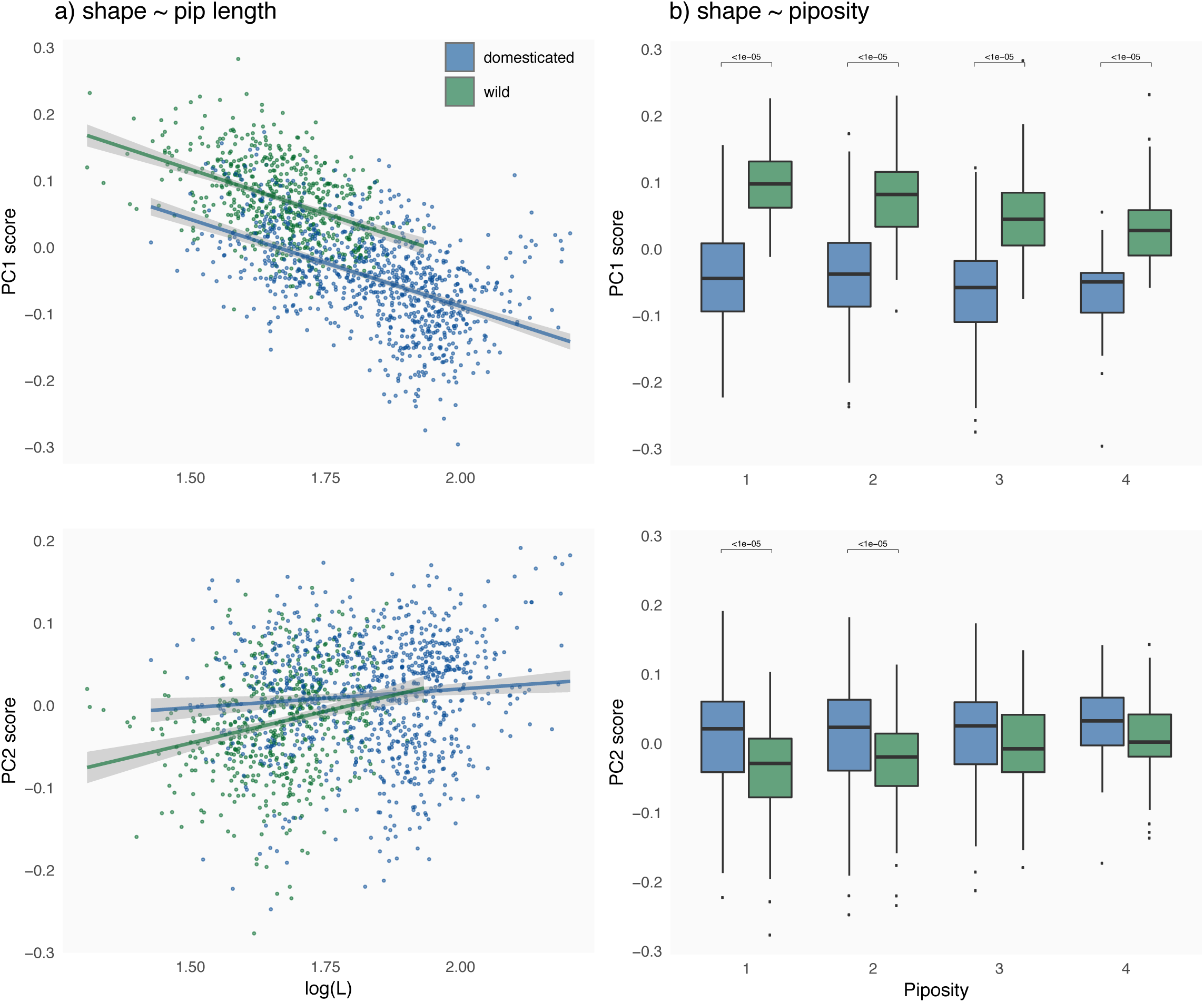
a) Regressions PC1 and PC2 versus pip length for domesticated pips (blue) and cultivation (green); b) boxplots for each piposity level for domesticated pips (blue) and cultivation (green). Differences are tested using Wilcoxon rank tests, and differences of P<10^− 5^, are indicated by brackets above graphs.

Regarding shape versus pip dimensions, pip_Length_, correlated to all other measurements, is itself correlated with position on PC1. Two regressions were justified (Analysis of covariance: df=1, F=362.7, P<10^−16^); their slope are identical (df=1, F=0.037, P=0.848) but their intercept differed between wild and domesticated. These two regressions were significant yet r^2^ were low (wild: P<10^−16^, adj. r^2^=0.195; domesticated: P<10^−16^, adj. r^2^=0.240 – Figure 5a). When PC1 and PC2 are considered jointly, two regressions were not justified (P=0.04) and the r^2^ was lower (P=0.04, adj. r^2^=0.181 – Figure 5a). The longer the pip is, the more “domesticate” it looks, particularly in terms of stalk prominence.

As concerns shape versus piposity, the latter is associated with shape changes on PC1 between wild and domesticated both overall (see above) and within levels (Wilcoxon rank tests, all P<10^−10^ – Figure 5b). Within domesticated accessions, differences were never significant. Within wild accessions, differences were not found between pairs of successive piposity levels but those between 1-3, 2-3 and 2-4 (all with P<10^−8^ – not shown). For PC2, general differences observed between domesticated and wild vanished for high piposity (1-pip: P<10^−12^; 2-pips: P<10^−9^; 3-pips: P=0.016; 4-pips: P=0.035 – Figure 5b). No differences within wild/domesticated and between successive piposities were found significant.

Mean shapes (Figure 6) illustrate these results. The mean absolute difference (MD) confirms that larger changes between extreme piposities are observed within wild grapevines (particularly for cultivated ones) and reveals that most of these changes affect the dorsal side of the pips (Figure 6).

**Figure 6.**
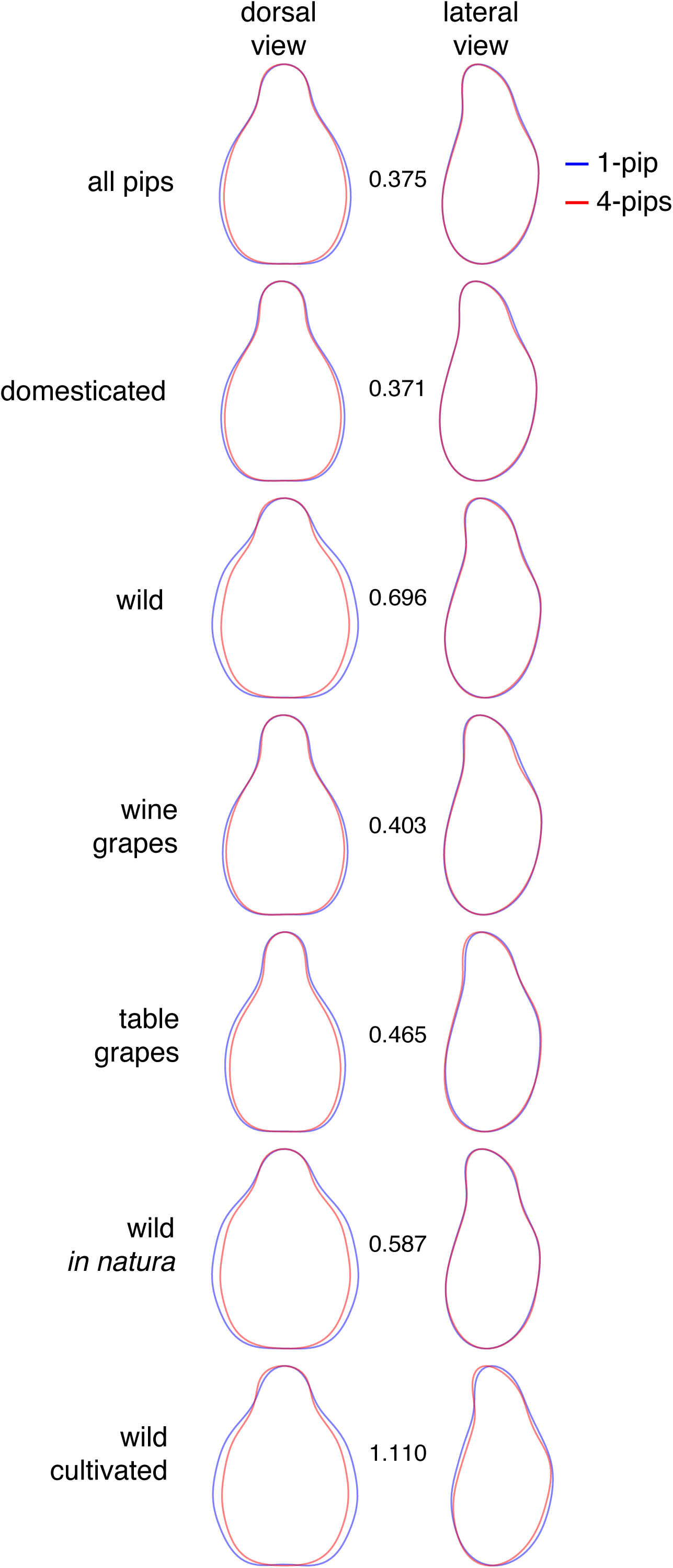
Mean shapes calculated for all pips sampled from berry with 1 (blue) or 4 pips (red), for different subsets (in rows) and for the dorsal and lateral views (in columns). Between the two views, an index of shape differences between these two extreme piposity levels (0=identical shapes; unit=as much differences as between wild and domesticated average shapes).

### Pip shape and size in relation to status, accession and piposity; consequences for archaeobotanical inference

The respective contributions of berry height, accession and piposity on the shape of pips (Figure 7) show that the accession is the factor affecting the most the pip shape. Among the different subsets, the accession factor has a higher impact on domesticated grapevine than on wild, and on cultivated wild accessions than on those collected *in natura*. By contrast, its contributions for wine and table domesticated varieties were similar. Here again, piposity and berry height both affect the pip shape of wild accessions but have a limited (piposity) and very limited (berry height) contribution for domesticated accessions.

**Figure 7.**
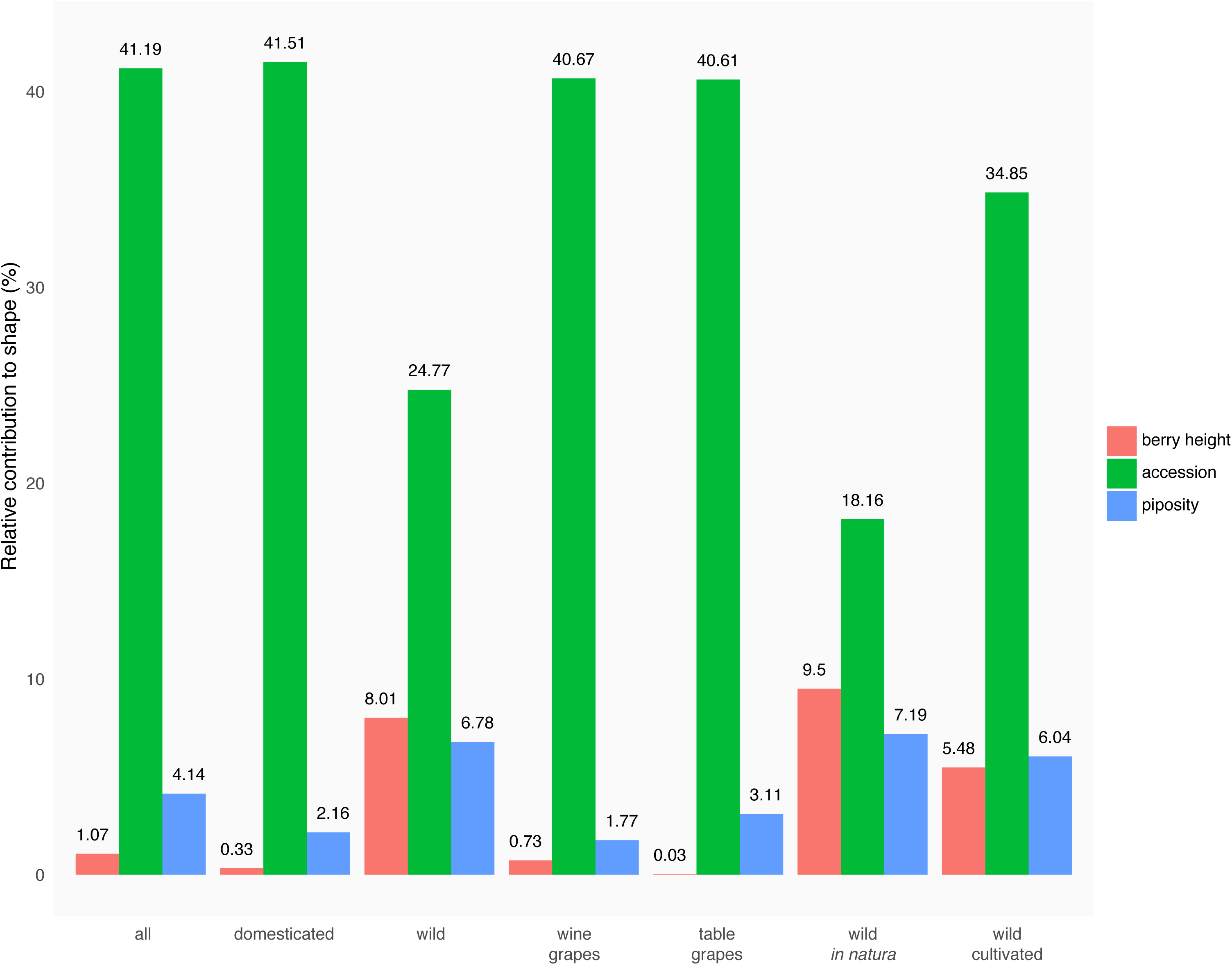
Relative contribution of berry height, accession and number of pips per berry (coloured bars) onto the shape of pips for different subsets.

Classification accuracies were compared using different training data and on different subsets (Figure 8). When different piposity levels were pooled, mirroring archaeobotanical admixtures, classification was very good at the status level (Figure 8a). Size + shape performed better (95%), than shape (93.7%) and size (92.5%) alone. When these models were evaluated on piposity subsets, they all have an accuracy above 91%, except for 4-pips berries. In all cases, accuracies were much higher than what could be obtained by chance alone. As expected, accuracies were lower at the accession level (Figure 8b) and when piposity levels were pooled, size + shape (89.8%) outperformed shape alone (81.3%) and size alone (46.3%). The same model ranking was observed on piposity subset, except for 4-pips berries. Overall, accuracies were nevertheless much better than chance alone.

**Figure 8.**
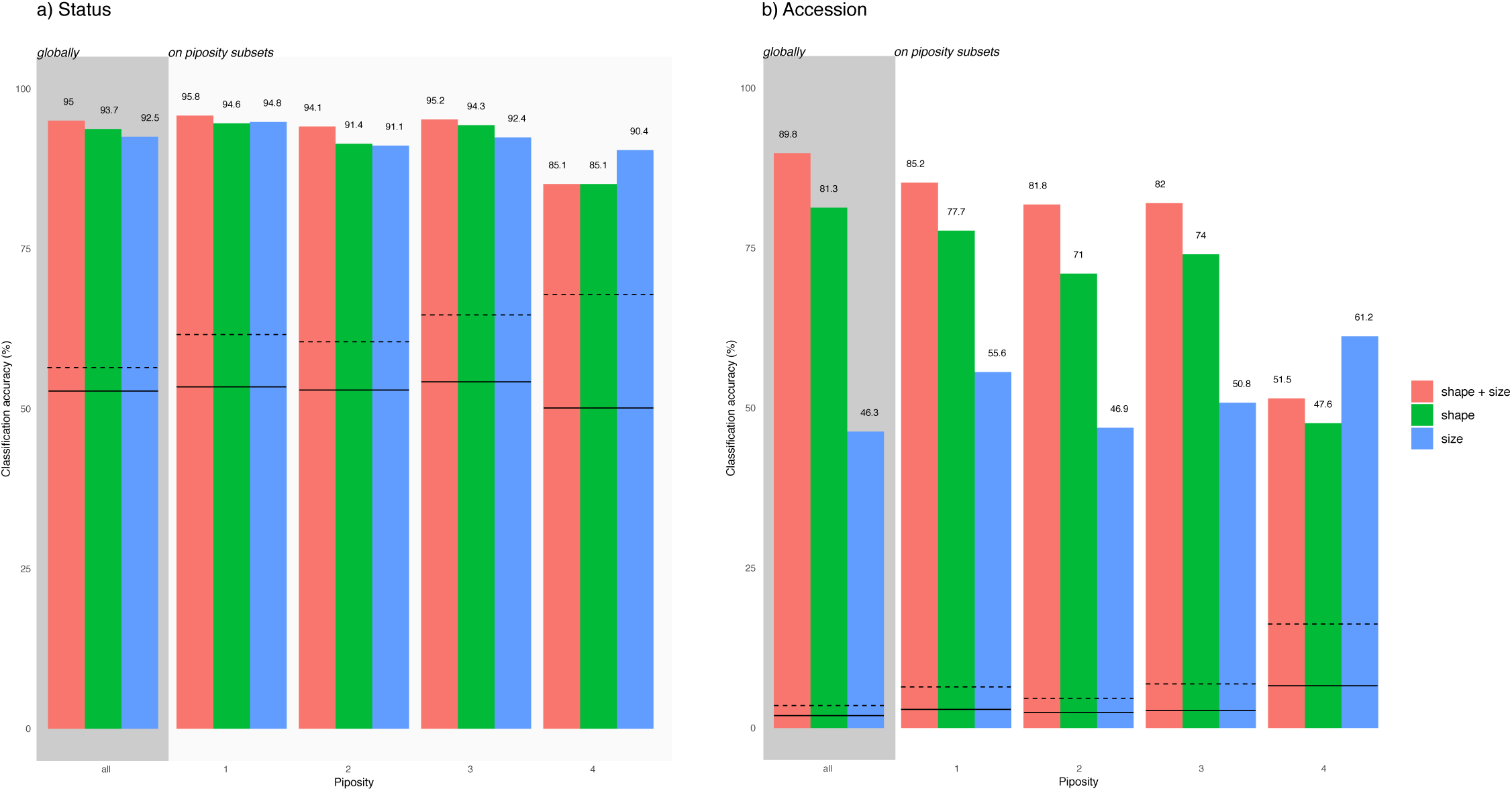
Classification accuracy (LDA leave-one-out) at the a) status and b) accessions levels. Models are trained (and evaluated) on the admixture of pips, then evaluated on piposity subsets. Different combinations of training data are used (Fourier coefficients of shape, lengths/mass measurements, both). Lines provide a random baseline and summarise 10000 permutations: solid line correspond to mean accuracy; dashed line to the maximal values obtained.

### Application to archaeological pips: can we infer the dimensions of the (vanished) berry dimensions from the (recovered) pips?

On modern material, we used the size of pips to predict berry heights and diameters. Both regressions show a significant interaction of the domestication status (berry_Diameter_: df=1, F=8369, P<10^−16^; berry_Diameter_: df=1, F=7730, P<10^−16^), and two regressions for berry diameter and two others for its height were obtained (Figure 9). All were significant (all P<10^−16^) yet the adjusted r^2^ were quite low (berry_Diameter_ adj. r^2^_wild_=0.585, adj. r^2^_domesticated_=0.491; berry_Height_ adj. r^2^_wild_=0.615, r^2^_domesticated_=0.511). Final models all used pip_Length_, pip_Thickness_, and at least one PC. Table 2). On unlogged (to compare “real” deviations obtained) berry diameter and height, the relative deviations were obtained (Figure C – ESM). Mean relative deviation per accession for berry_Diameter_ ranged from −12.9% to +10.3% for wild, and from −22.9% to +17.7% for domesticated; for berry_Height_ they ranged from −13.0% to +13.1% for wild, and from −29.4% to +29.4% for domesticated. The average predictions were all centred (on zero) ±1.6%.

**Table 1.**
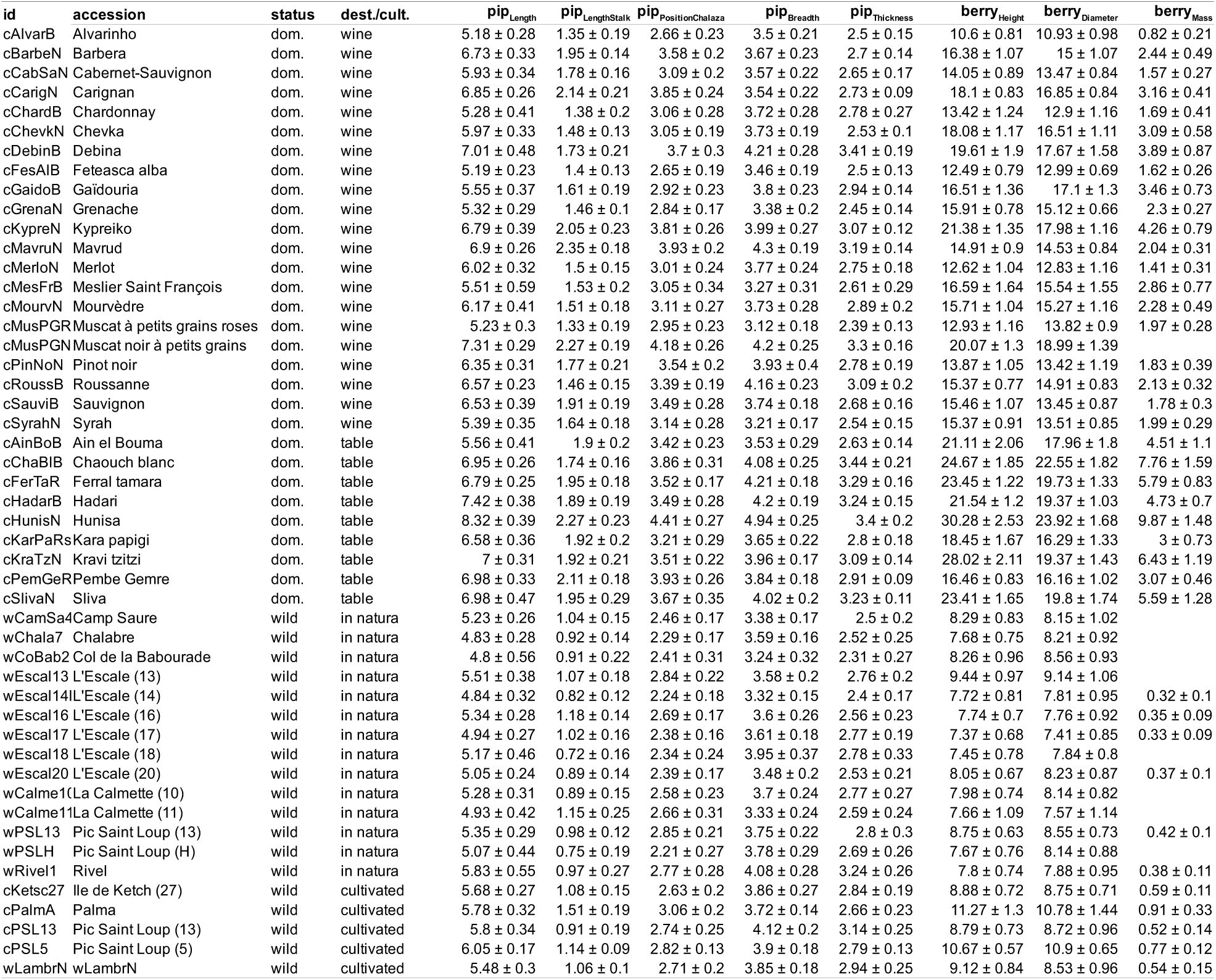
Accessions used in this study. D: domesticated; W: wild; Wn: wine grape; Tb: table grape. For domesticated grapevines, names correspond to the variety names. Dimensions are reported with mean±sd and given in mm, except for berry_mass_ which is expressed in g.

**Table 2.** Estimates for pip lengths used to infer berry dimensions. Variables were all logged; so that berry_Diameter_ (in mm and for wild) can be obtained with exp[0.65163 +1.19914×log(pip_Length_) +0.10617×log(pip_PositionChalaza_) −0.13263×log(pip_Breadth_) - 0.45449×log(pip_Thickness_)].

|  |  | Estimates of predictors |  |  |  |  |  |  |  |
| --- | --- | --- | --- | --- | --- | --- | --- | --- | --- |
| log Predicted: | subset | Intercept | log pip<sub>length</sub> | log pip<sub>LengthStalk</sub> | log pip<sub>PositionChalaza</sub> | log pip<sub>Breadth</sub> | log pip<sub>Thickness</sub> | PC1 | PC2 |
| berry<sub>Diameter</sub> | wild | 0.689 | 1.136 | — | — | — | -0.438 | -0.223 | 0.209 |
|  | domesticated | 1.298 | 0.478 | — | 0.149 | -0.150 | 0.520 | — | 0.430 |
| berry<sub>Height</sub> | wild | 0.838 | 0.732 | — | 0.216 | 0.171 | -0.318 | -0.524 | 0.258 |
|  | domesticated | 0.760 | 1.312 | 0.279 | — | -0.783 | 0.584 | 1.021 | — |

**Figure 9.**
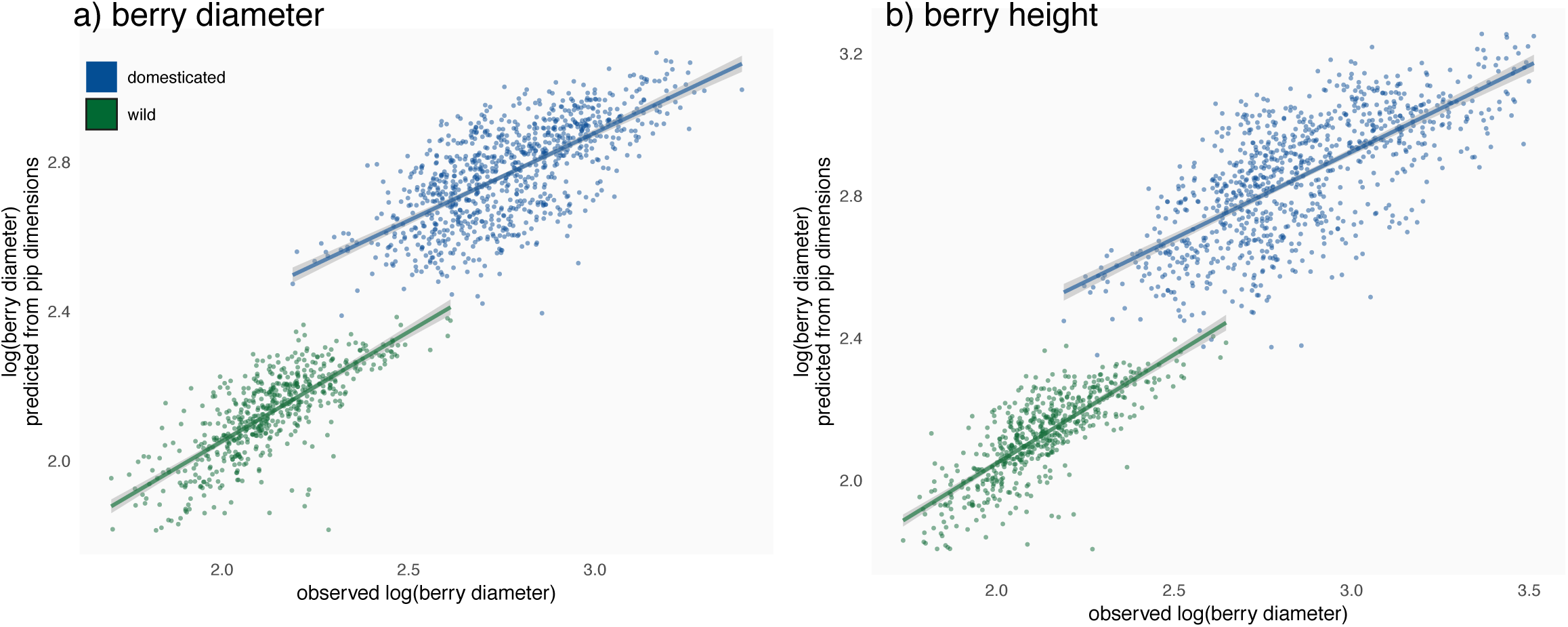
Regressions for berry dimensions from pip dimensions, obtained on modern material: predicted versus actual (logged) berry dimensions at the domestication status level (all accessions). Columns are for berry diameter and height, respectively.

Then, these four models were applied on the archaeobotanical material after being classified at the wild/domesticated level using LDA. 46 pips (22%) were classified with a posterior probability <0.8 and were filtered out. Among the remaining pips, 114 (72%) were classified as domesticated and 45 (28%) as wild. When compared to their modern analogues (Figure 10), the length of “domesticated pips” were closer to those of wine varieties than table varieties; the lengths of “wild pips” were intermediate between wild accessions collected in their habitat and those cultivated. For archaeological pips identified as domesticated, both inferred berry height and diameter were intermediate between wine and table modern varieties yet closer to wine ones. Similarly, for wild archaeological material, inferred berry height and diameter were intermediate between wild collected in their habitat and those grown in collection but closer to the former.

**Figure 10.**
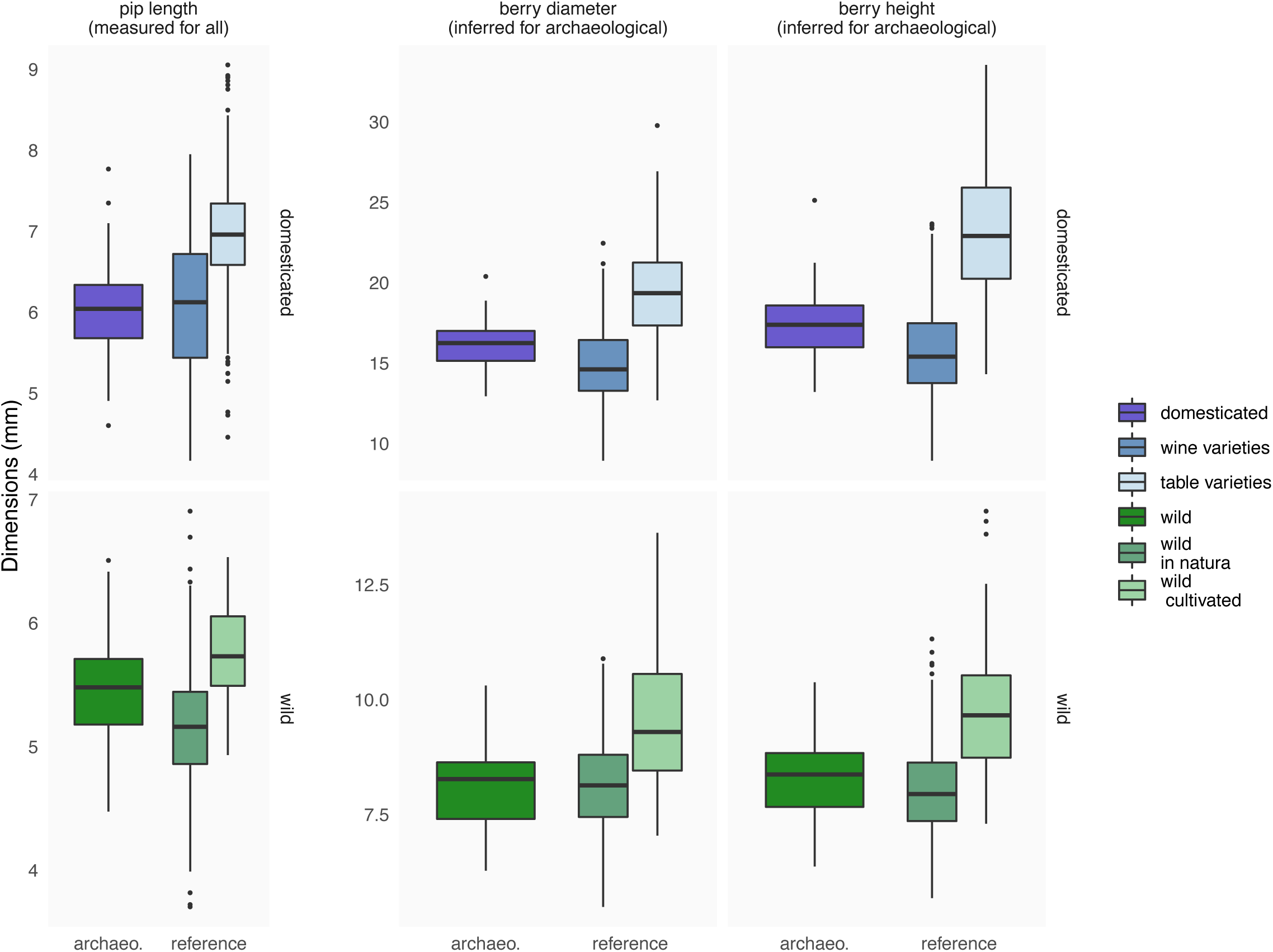
Distribution of pip lengths (observed for all), berry height and diameter inferred from pip dimensions for the archaeological material from Sauvian – La Lesse. Rows distinguish domesticated and wild grapes since separate regressions were required.

## Discussion

This study opens major fronts in our understanding of *Vitis vinifera* phenotypic changes under domestication and helps disentangle the interplay of the number of pips per berry, berry dimensions, domestication, pip shape, varietal diversity and cultivation practices in both wild and domesticated grapevines. We discuss implications for *Vitis vinifera* eco-evo-devo and perspectives for archaeobotanical studies for which a possible application is proposed.

### Patterns of covariation between the form of the pip, the form of the berry and the piposity

With two ovules per ovary, and two ovaries per berry, the theoretical maximum number of pips per berry is four, yet one was observed with five. Such abnormal piposity have been reported, for berries having more than two ovaries [8,36]. Most berries had two pips, and more than 70% had only one or two. This is in accordance with previous publication [35]. There were no differences neither between domesticated or wild (Figure 2), nor between cultivated wild individuals and those collected in their habitat. Higher piposity could be expected for domesticated grapevine (since they have hermaphroditic flowers), and more generally for cultivated accessions (since the pollen rain is expected to be lower, or even limiting, in the natural habitat).

Overall, and this is no surprise, wild pips and berries are smaller than their domesticated counterparts. Similarly, pips and berries of wine varieties are smaller than those of table varieties, as pips and berries of wild grapevines collected in their habitat are smaller than those from cultivated wild individuals. This study details the effect of piposity on the pip form reported by previous studies (Bouby et al., 2013; Houel et al., 2013; Negrul, 1960; Olmo, 1995). Among vertebrate dispersed plants, the reward (the fruit pulp mass) associated with a given seed mass is commensurate with work required to move it, and is expected to scale relatively [37]. For wild grapevine, and vertebrate dispersed plant species in general, berry and pip dimensions/masses are expected to be constrained by their dispersers and by a general trade-off between pip size and number [38].

For all but wine varieties, the higher the piposity the longer the pip and the bigger the berry in which they develop (Figure 3). For these groups, it seems that more numerous pips are not limited by space or nutrients but rather contribute the development of bigger berries. The stages of berry development are well known [39,40] and can be divided into two phases of enlargement. The first, prior to anthesis, is a period of rapid berry growth mostly due to cell division. After anthesis, berry growth is largely due to cell enlargement and it has been suggested that pip growth may also increases cell mitosis in the developing berry (Ojeda et al., 1999). Auxins, cytokinins and gibberelins, upregulated shortly after fertilisation in grapevine ovaries, are likely to trigger berry growth by cell expansion [35].

The absence of positive (or even negative) correlations between piposity, pip and berry dimensions for wine varieties remains unclear. For these varieties, the regulation, if any, may be at the bunch or stock scale, whether it has been selected (for example to concentrate sugars, aromas and flavours) or it is a by-product of another trait under selection. Since table varieties are larger than wine varieties, the berry dimensions of the latter cannot be argued to have reached a developmental limit.

Finally, bivariate correlations concerning berry dimensions and mass are the strongest observed. This indicate robust allometries between berry size and mass, in other words that berry largely remain ellipsoid in shape, independently of their dimensions.

### Morphometrics and domestication as a wedge into grapevine eco-evo-devo

For grapevine and domesticated plants in general, domestication results in a change of desirable phenotypic patterns (bigger fruits for instance) but also releases many “natural” constraints such as dispersion [42]. Cultivation practices such as pruning may explain why wild individuals grown in collection have bigger berries for higher piposity: the number of bunches is reduced, leading to larger pips. Cultivation also reduces growth constraints such as competition for water and light, self-supporting and climbing costs, those related to dispersers, etc.

Evidence of plastic and canalized phenotypic expression may be fuel for further eco-evo-devo studies. The latter brings a conceptual and experimental framework that relies on environmentally mediated regulatory systems to better understand ecological and evolutionary changes [43]. Here, the norm of reaction of the pip size and shape, along increasing piposity and berry dimensions, is clearly different at the three investigated levels: between wild and domesticated, between wine and table, between wild individuals grown in collection and those collected in their natural habitat.

### Consequences for archaeological inference

Taken independently or in pairwise comparisons, some pip lengths differences between wild and domesticated appear more “robust” to increasing piposity, notably pip_LengthStalk_ and pip_PositionChalaza_ (Figure 3a; Figure A Supplementary information). Their interest in discriminating domestication status has long been used, including when archaeological material is charred (Bouby et al., 2018; Mangafa & Kotsakis, 1996; Smith & Jones, 1990).

Overall, shape variability tends to distinguish the wild and domesticated grapevines (Bouby et al., 2013; Terral et al., 2010). Similar to the positive correlation between piposity and pip length of wild accessions, piposity is also correlated with shape changes for wild accessions (Figure 5). Most of these changes affect the dorsal side of wild pips, particularly when piposity is high. There is as much as ∼70% difference, between 1- and 4-pips for wild accessions, than those observed between average wild and domesticated. Differences in extreme piposity are even larger than this “domestication gap”, for the cultivated wild. This does not answer the question whether past vineyards cultivated “true” wild grapevines or “weakly” domesticated forms (Bouby et al., 2013; Pagnoux et al., 2015) but it points out how piposity and cultivation practices contribute to this confusion, further enhancing the continuum of pip forms.

Pip shape being largely used in archaeobotany, it was crucial to point out which factors contribute to its variability, or at least covary with it, and if they could preclude identification of archaeological remains. Here, the main factor associated to pip differences was, by far, the accession and it was even more important for domesticated accessions; in other words accession effect appears stronger than domestication (Figure 7). Relatively to accession, berry height and piposity poorly contributed to observed differences. This confirms the usefulness of shape and its robustness to identify morphotypes that are shape varietal archetypes. It may also indicate that domestication favours pip shape diversification whether this results from genetical linkage with selected loci or is the product of drift.

Here, we show the reliability of classification, independently of piposity. Indeed, classification accuracies at the status level were all high (Figure 8), even when the models trained on the pip admixture where evaluated on piposity subsets. Shape was nonetheless superior to size alone in discrimination power but, when considered jointly, the classification was improved. Whenever possible, size should thus be included along morphometric coefficients and used jointly in classification models. Accuracies at the accession levels evidenced even more clearly the latter conclusions.

Shape is overall more robust than sizes when models were evaluated on piposity subsets. The only exception, for both status and accession levels, were obtained on the 4-pips subset. Our experimental design reflects real-world admixtures, and sample sizes of all studied factors were not balanced. That being said, results here evidence that such bias in the piposity structure is very unlikely to affect archaeobotanical identification either at the status or accession levels.

### An application on archaeological material: inferring berry size from pip

Berry is very likely home to the most selected traits, from the beginnings of domestication to varietal breeding and diversification times. Unfortunately, its dimensions cannot be quantified directly on archaeological material where fleshy parts are usually absent or too degraded. The only route to investigate changes in berry shape and size is through actualistic inference based on pip, and trained on modern material. Here, multiple regressions on pip dimensions show that berry diameter (Figure 9a) and height (Figure 9b) were not perfectly predicted but nevertheless centred on zero and overall in the ±25% range.

Our archaeological application used material from Sauvian - La Lesse, a Roman farming establishment involved in wine production, were an admixture of wild and domesticated type is attested (Figueiral et al., 2015). As in many cases in Southern France, the presence of numerous wild type pips in a vinicultural site let us consider that these vines were locally cultivated to make wine in Roman times. Berry dimensions inferred from pips of this site are intermediate between the wild growing in their habitat and those cultivated (Figure 10). This may suggest that wild, or weakly domesticated, individuals were cultivated in Roman vineyards. The berry dimensions inferred for domesticated varieties were closer to modern wine varieties than to table ones. This is congruent with the wine production attested at this period and in this region (Figueiral et al., 2010; Figueiral et al., 2015).

## Conclusion

The main finding of this systematic exploration of berry and pip form covariation is that for wild grapevine, the higher the piposity, the bigger is the berry and the longer is the pip. For both wild and domesticated, the longer is the pip, the more its shape looks like “domesticated”. Further studies will clarify the contribution of cultivation practices contribution on pip shape, largely used in archaeobotanical studies to better understand viticulture history. These findings pave the way for dedicated studies to shed light on genetic, functional and evolutionary changes that occurred in *Vitis vinifera* between the pip, its reproductive unit, and the berry, its dispersal reward and the main target of its domestication and varietal improvement.

## Material and Methods

### Statistical environment

Statistical analyses were performed using the R 3.6.2 environment [46], the package Momocs 1.3.0 for everything morphometrics (Bonhomme et al., 2014) and the tidyverse 1.2.1 packages for data manipulation and most graphics [48]. Alpha significance level was chosen equal to 10^− 5^ all along analyses. This level both ensure marked differences for subsequent archaeobotanical application and an overall alpha level below 0.05 when repeated tests were done (i.e., a Bonferroni correction).

### Nomenclature

Hereafter, *status* designates compartment (domesticated vs. wild); *accession* designates the variety (or cultivar, or *cépage*) for domesticated grapevine and the individual for wild grapevine; synecdochically, a domesticated/wild pip/berry refers to the accession they were collected from; *cultivation* designs whether wild individuals were *cultivated* (grown in field collection) or sampled *in natura*; *form* is used when *shape* and *size* are used in combination; “piposity” is short for “given a pip, the number of pips in the berry where it was sampled”.

### Modern and archaeological material

The modern reference material included 49 accessions (30 domesticated and 19 wild) from Euro-Mediterranean traditional cultivars and wild grapevines (Table 1). Fourteen wild grapevines were collected at ripeness in their habitat, and five were cultivated in the French central ampelographic collection (INRA, Vassal-Montpellier Grapevine Biological Resources Center; https://www6.montpellier.inra.fr/vassal), along with the domesticated accessions. Of the domesticated accessions, 21 were wine varieties and 9 table varieties. For each accession, 30 normally developed berries have been haphazardly collected from a single, fully ripe bunch.

Archaeobotanical material comes from two wells at the Roman farm of Sauvian - La Lesse, extensively described elsewhere (US3022, US3063, US3171 and US3183 in Figueiral et al., 2015). These archaeological layers were dated to 2025-1725 BP based on pottery and coins. The waterlogged conditions ensured very good preservation the pips used in this study (N=205).

### Traditional measurements

On modern material, the berry diameter (berry_Diameter_), height (berry_Height_) and mass (berry_Mass_) were obtained before dissection (Table 1). Mass was not available for 9 accessions that were removed from further analyses involving mass. Then, the number of pips (hereafter “piposity”) was recorded and one pip was randomly chosen. A single berry from the variety “Kravi tzitzi” was found with 5 pips and was discarded from further analyses. The final dataset thus consisted of 1469 pips (48 accessions × 30 pips + 1×29).

All pips, archaeological and modern, were photographed in dorsal and lateral views by the same operator (TP) using an Olympus SZ-ET stereomicroscope and an Olympus DP camera. On each pip, five length measurements were manually recorded by the same operator (LB) and using ImageJ (Rasband, 2008, Table 1, Figure 1): total length (pip_Length_), length of stalk (pip_LengthStalk_), position of the chalaza (pip_PositionChalaza_), breadth (pip_Breadth_) and thickness (pip_Thickness_). All length measurements were log-transformed to focus on relative changes and minimize size differences; the mass was log cubic-root transformed for the same reason (Bouby et al., 2013).

As preliminary analyses on modern material, differences between average piposity were tested using generalized linear model with Poisson error; differences in their distributions were tested using two-sided Fisher’s exact tests on count data.

### Testing the covariation between pip and berry size in relation to the number of pips

On modern material, three sets of differences in pips and berries measurements were tested using multivariate analysis of covariance: i) the interaction between status and piposity; ii) if the latter was significant, we also tested differences between status for a given piposity level; iii) whether the average piposity differs between status. These three possible sets of differences were tested between different subsets: domesticated and wild accessions; wine and table varieties for domesticated accessions; cultivated wild individuals and those collected *in natura*. Piposity was then discarded and sets were compared using Wilcoxon rank tests.

Bivariate comparisons were explored between the domesticated and wild accessions (discarding piposity), and tested with an analysis of covariance. When the domestication status was significant, separate regressions were tested and, if significant, the adjusted r^2^ was obtained.

### Testing the covariation between pip and berry shapes in relation to the piposity

For pips, shapes data were extracted from the dorsal and lateral outlines. 2D coordinates were extracted from photographs, centred, scaled, aligned along their longer axis (using the variance-covariance matrix of their coordinates) and normalized for the position of their first point before elliptical Fourier transforms (EFT). These preliminary steps removed positional, size, rotation and phasing differences between outlines and EFT could then be used without numerical normalization [50]. EFT were performed on the dorsal and lateral views separately, and the number of harmonics was chosen to gather 99% of the total harmonic power (8 for both views). This generated 64 coefficients for each pip (2 views × 8 harmonics per view × 4 coefficients per harmonic).

To explore the overall variability of shape, a principal component analysis (PCA) was calculated on the full matrix of coefficients. The first two PCs (see Results) were used as synthetic shape variables. To test the effect of piposity and pip dimension on pip shape, the same approach than for length measurements using PC1 and PC2 as the response variables. To test the relation between shape and pip length (only pip_Length_ was used), analyses of covariance first tested if separate regressions were justified. Then Wilcoxon tests were used to test for shape differences between and within piposity levels.

To visualize shape differences between extreme piposity levels (1 and 4), mean shapes for the dorsal and lateral views were calculated on the matrix of coefficients. These differences were quantified with the mean absolute difference (MD) between each sets of Fourier coefficients. To make these differences meaningful, they were divided by the mean difference of Fourier coefficients between cultivated and wild accessions with all piposity levels pooled. For each subset, MD was calculated as: (|coefficients_subset, 1-pip_ - coefficients_subset, 4— pips_ |)/(| coefficients_domesticated, all pips_ - coefficients_wild, all pips_ |). For example, a MD equals to 0 would indicate no difference between pips with a piposity of 1 or 4; a MD greater than unit would indicate more differences relatively to differences that exist between domesticated and wild individuals.

### Pip shape and size in relation to status, accession and piposity

To quantify the respective contribution of berry dimensions, accession and piposity onto pip shape, a multivariate analysis of variance used the following model: all Fourier coefficients ∼ berry_Height_ + accessions + piposity within accession. Since it was highly correlated to other berry measurements (see Results), only berry_Height_ was used to describe berry dimensions. The contribution of each variable is the ratio of its sum of squares over the total sum of squares (including residuals). Again, this is tested on the different subsets of interest (Figure 7).

Linear discriminant analyses (LDA) were used to evaluate whether piposity could preclude status and accession classification accuracies. Different combinations of predictors (sizes; shape; sizes + shape) were evaluated to benchmark their performance to classify the pips to their correct status and accession (Figure 8).

Given a combination of (status, accession) × (sizes+shape, sizes, shape), a leave-one-out cross-validation was used to assess classification accuracies, evaluated on all pips, to mirror archaeological admixtures where piposity is unknown (Figure 8). To cope with unbalanced group structures, we calculated a baseline for each subset that estimates the mean and maximum accuracy one can obtain by chance, using 10^4^ permutations (see (Evin *et al*., 2013). If the accuracy observed is higher than the maximum value obtained using permutations, the LDA can be considered to perform better than random, with an estimated alpha below 10^−4^.

### Predicting the dimensions of the archaeological berry dimensions

Separate multivariate regressions were calculated on the modern material, for berry height and diameter (using the five length measurements on pips). As predictor variable, we used length measurements (for dimensions) and the first two principal components (for shape). The difference between domesticated and wild grapevines regressions was first tested using an analysis of covariance: two regressions (one for cultivated, one for wild) were obtained for the berry height and two others for its diameter (Figure 9). These four regressions were fitted using stepwise regression with backward elimination based on the AIC (Venables & Ripley, 2002), and started with full models: berry_Height/Diameter_ for wild/domesticated ∼ pip_Length_ + pip_LengthStalk_ + pip_PositionChalaza_ + pip_Breadth_ + pip_Thickness_ + PC1 + PC2, all but PCs were log-transformed). Then, archaeological pips were classified into domesticated or wild using an LDA trained using the same variables but of modern pips. Pips assigned to wild/domesticated with a posterior probability <0.8 were filtered out. Finally, the berry height and diameter of this archaeological material were inferred using the corresponding models (Figure 10).

## Acknowledgements

This study is funded by ANR project “Vignes et vins en France du Néolithique au Moyen Âge. Approche intégrée en archéosciences” (PI: Laurent Bouby) and supported by the OSU-OREME (https://oreme.org/) that helped to the constitution of the wild grape pip collection. We are grateful to the INRA Vassal-Montpellier grapevine collection (Marseillan-Plage, France) that provided all the pips from cultivated varieties and cultivated wild grapes. We warmly acknowledge Michael Wallace for his help with English.

## Author contribution

LB, SP, SI and JFT conceived the ideas and designed methodology; SP, SI, TP, IF and LB collected the data; VB, led analysed the data with important contributions from SP, AE, JFT, LB, TL and RB; VB led writing of the manuscript with the help of all co-authors, and chiefly SP.

---

**Figure A (ESM).**
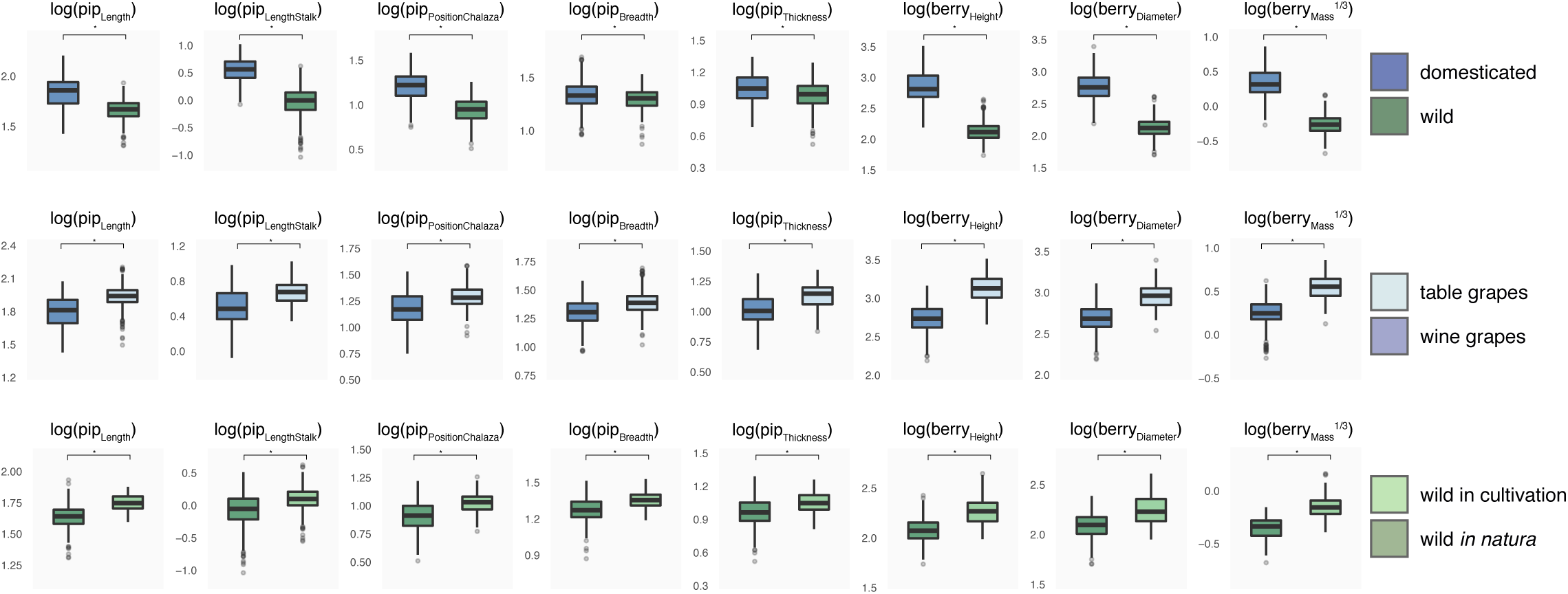
Comparisons of (logged) lengths and (cubic-rooted) mass measurements. On rows are displayed different subsets: a) wild and domesticated grapevines, b) for domesticated accessions, table and wine varieties and, c) for wild accessions, those collected *in natura* and others cultivated as domesticated varieties. Different piposity levels are pooled (see Figure 3 for the detail). Differences are tested using Wilcoxon rank tests and all of them have a P<10^−5^.

**Figure B (ESM).**
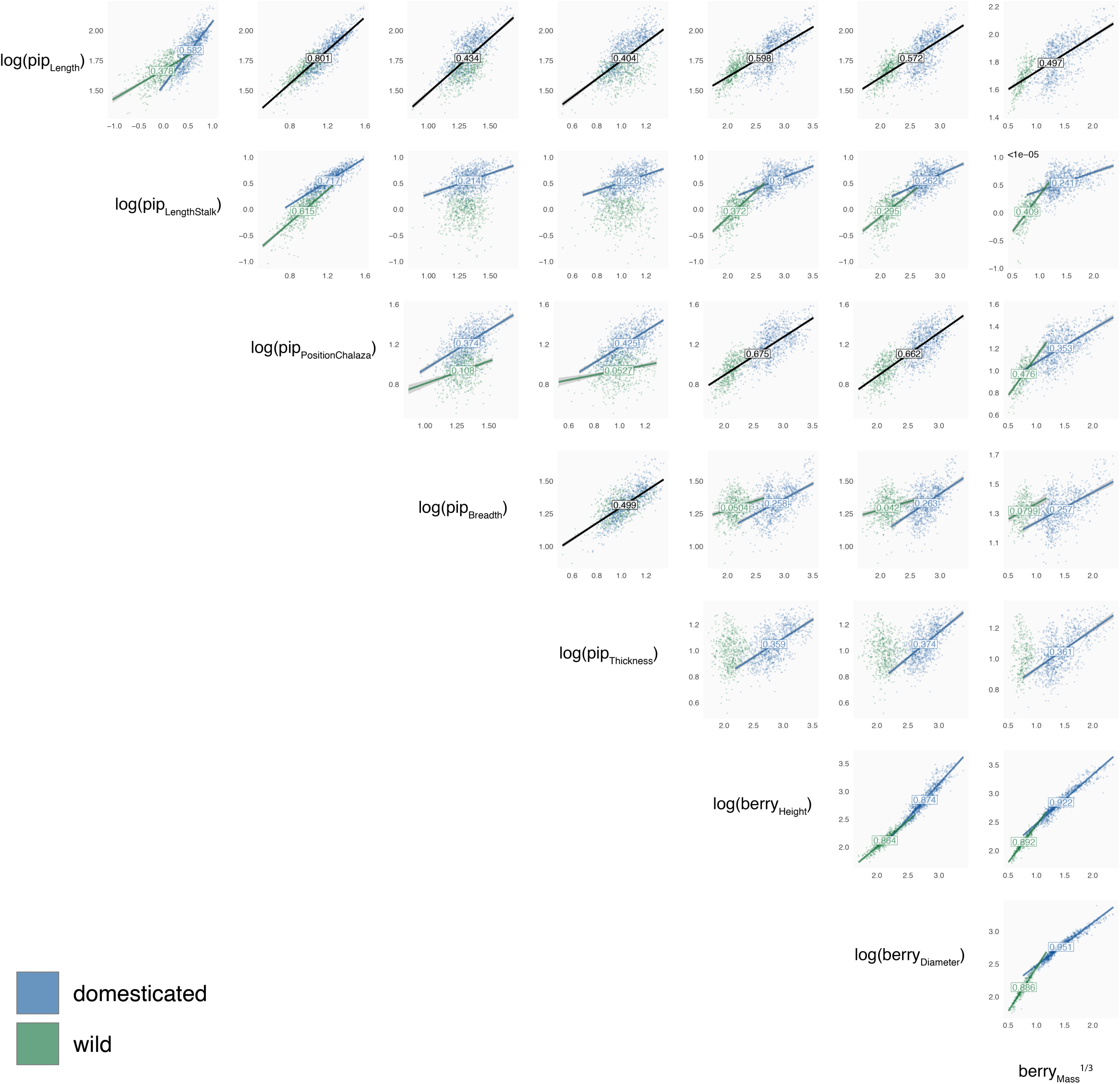
Bivariate pairwise plot between (logged) lengths and (cubic-rooted) mass measurements. For the sake of readability, only the wild versus domesticated status are displayed using different colours (green for wild; blue for domesticated). If two regressions are justified, then they are shown using the corresponding colours; otherwise a single regression line is showed in black. Then, for each regression, the correlations are tested and, if significant, the adjusted R^2^ is displayed on the regression lines.

**Figure C (ESM).**
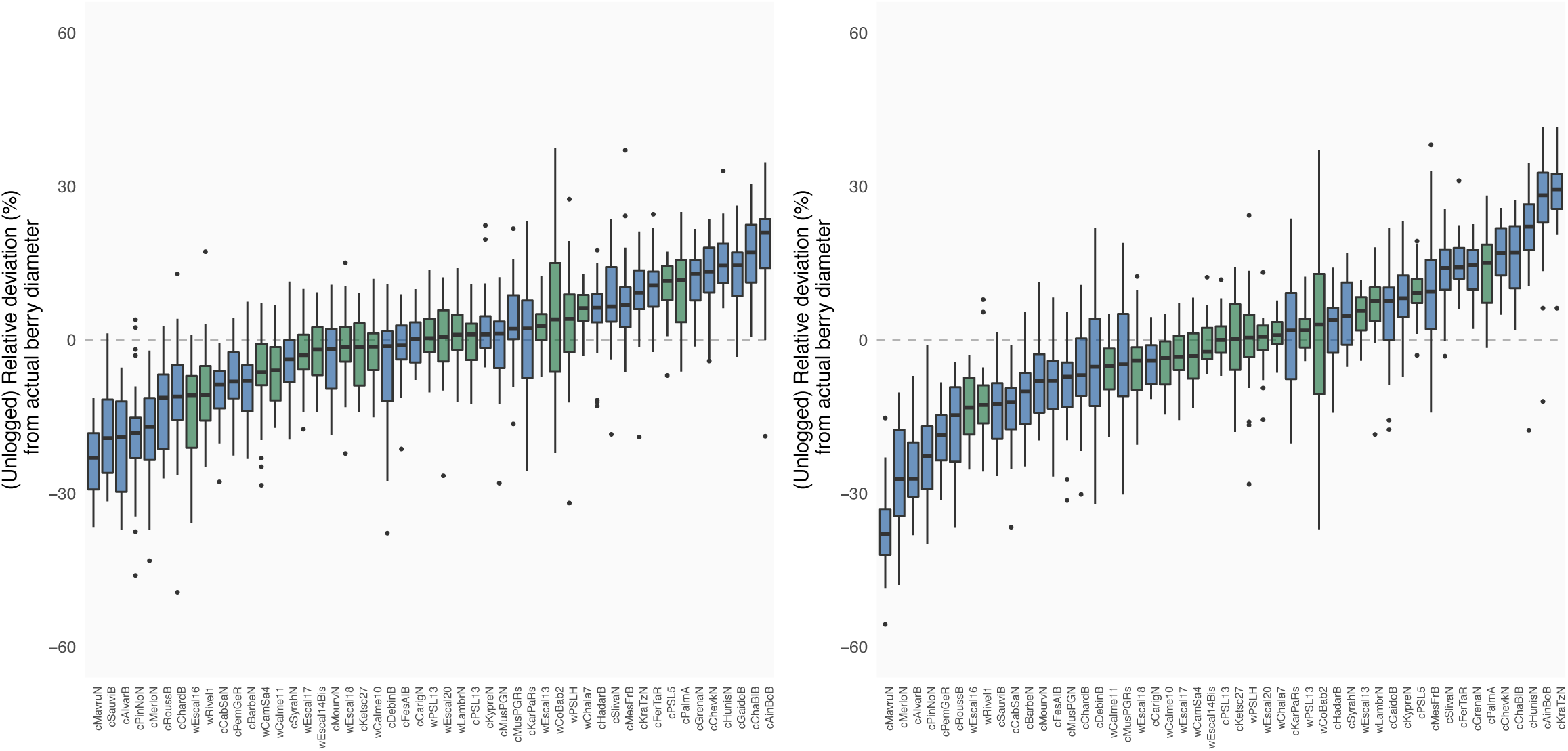
Predictions obtained from regressions for berry dimensions from pip dimensions on modern material. The relative deviation, at the accession level, and for unlogged measurements. Columns are for berry diameter and height, respectively.

